# Induced and primed defence responses of *Fragaria vesca* to *Botrytis cinerea* infection

**DOI:** 10.1101/692491

**Authors:** Raghuram Badmi, Yupeng Zhang, Torstein Tengs, May Bente Brurberg, Paal Krokene, Carl Gunnar Fossdal, Timo Hytönen, Tage Thorstensen

## Abstract

Strawberry is a high-value crop that suffers huge losses from diseases such as grey mould caused by the necrotrophic fungal pathogen *Botrytis cinerea*. Pesticides are heavily used to protect the strawberry crop, which raises environmental and human health concerns and promotes the evolution of pesticide resistant strains. Upregulating or priming the plants’ defences may be a more environmentally sustainable way of increasing disease resistance. Using *Fragaria vesca* as a model for the commercially grown octaploid strawberry *Fragaria × ananassa*, we investigated the transcriptional reprogramming of strawberry upon *B. cinerea* infection and the effectiveness of four priming chemicals in protecting strawberry against grey mould. First, we found that the transcriptional reprogramming of strawberry upon *B. cinerea* infection overlapped substantially with the transcriptome responses induced by *Phytophthora cactorum* (Toljamo et al., 2016), including the genes involved in jasmonic acid (JA), salicylic acid (SA), ethylene (ET) and terpenoid pathways. Furthermore, we investigated the effectiveness of previously identified priming chemicals in protecting strawberry against *B. cinerea*. The level of upregulated or primed resistance depended on the priming chemical itself (β-aminobutyric acid (BABA), methyl jasmonate (MeJ), *(R)*-β-homoserine (RBH), prohexadione-calcium (ProCa)) and the application method used (foliar spray, soil drench, seed treatment). Overall, RBH effectively primed strawberry defences against *B. cinerea*, whereas BABA and ProCa were not effective and MeJ showed mixed effects. Our results not only identify ways to effectively upregulate or prime strawberry defences against *B. cinerea*, but also provide novel insights about strawberry defences that may be applied in future crop protection schemes.

## 2. Introduction

Agricultural crops are vulnerable to attack by many pests and pathogens that kill plants, reduce yields, and thus cause huge economic losses. Fungal diseases alone are estimated to reduce crop yields by about 20% and cause 10% postharvest damage worldwide (Fisher *et al*., 2018). Fungicides (Fisher *et al*., 2018) and other pesticides (Gould, Brown and Kuzma, 2018) are the most frequently used crop protection strategy around the world. Pesticide use raises health concerns in humans, can harm the environment and can select for resistant pests and pathogens that ultimately may render these chemicals ineffective (Fisher *et al*., 2018; Gould, Brown and Kuzma, 2018). To increase the sustainability of agriculture there is a need to reduce pesticide use and develop more environmentally benign pest management solutions.

Plant defence priming agents that induce or prime plant resistance against diseases are promising alternatives for future pest management strategies. These defence priming agents can be applied to the plants as either a soil-drench, spray or seed treatments. One well-characterized defence priming chemical is β-aminobutyric acid (BABA). It has been shown to effectively prime plant defences against a broad spectrum of pathogens (Cohen, 2002; Wilkinson *et al*., 2018) by inducing both salicylic acid (SA) dependent and SA independent defences (Ton *et al*., 2005). Another well-known priming agent, methyl jasmonate (MeJ), has been shown to induce resistance in many plant species including strawberries (Saavedra *et al*., 2017) and grape berries (Wang *et al*., 2015) when applied directly onto the fruits. However, systemic application of BABA and MeJ can also suppress plant growth and reduce crop productivity (van Hulten *et al*., 2006; Yang *et al*., 2012). Recently, Buswell et al. (2018) identified a structural analogue of BABA, *(R)*-β-homoserine (RBH; R-3-amino-4-hydroxy butanoic acid), that induces broad spectrum resistance in *Arabidopsis thaliana* (Arabidopsis) and tomato (*Solanum lycopersicum*) without negative effects on plant growth.

Another potential priming agent is prohexadione-calcium (ProCa). This inhibitor of gibberellin biosynthesis has been shown to induce resistance against the bacterial disease fire blight in apple and pear (Yoder, Miller and Byers, 1999; Rademacher, 2004; McGrath *et al*., 2009), and against apple scab caused by the hemibiotrophic fungus *Venturia inaequalis* (Spinelli *et al*., 2010). The mechanism of ProCa-induced resistance is unknown, but is believed to involve biosynthesis of antimicrobial flavonoid compounds (Spinelli *et al*., 2005; McGrath *et al*., 2009).

Plant resistance can also be primed by treating seeds with defence priming agents before germination. For example, melon plants derived from seeds treated with acibenzolar-S-methyl (also called benzothiadiazole, BTH) and MeJ display induced resistance against different fungal plant pathogens (Buzi *et al*., 2004), and tomato plants derived from seeds treated with jasmonic acid (JA) are resistant to infection by *Botrytis cinerea* and herbivory (Worrall *et al*., 2012). However, seed priming may also have more complex effects. Treating tomato seeds with BABA primes resistance against powdery mildew but increases susceptibility to *B. cinerea* infection (Worrall *et al*., 2012).

The necrotrophic fungal pathogen *B. cinerea* has a very broad host range and causes grey mould disease in many different crop plants, including strawberry (Dean *et al*., 2012; van Kan, Shaw and Grant-Downton, 2014; Veloso and Kan, 2018). Due to its ability to infect more than a thousand plant species, its great economic importance and ease of transformation, *B. cinerea* is the most extensively studied necrotrophic fungus (Dean *et al*., 2012; Veloso and Kan, 2018). Infection by *B. cinerea* induces major transcriptional reprogramming in Arabidopsis. Nearly 10,000 genes are differentially expressed (Windram *et al*., 2012), including genes involved in major defence related signalling and biosynthesis pathways such as the salicylic acid (SA), ethylene (ET) and jasmonic acid (JA) pathways (Benito *et al*., 1998; Williamson *et al*., 2007). Most of these transcriptional changes occur within 48 hours of infection, before significant disease symptoms can be observed (Windram *et al*., 2012).

Commercial strawberry (*Fragaria × ananassa*) is a high-value crop that can suffer huge losses from pathogens like *B*. cinerea and it would therefore be of great value if strawberry defences could be primed. Different priming agents and methods have been demonstrated to increase disease resistance in strawberry. For example, harvested fruits immersed in a BABA solution showed increased resistance to *B. cinerea* infection in a concentration-dependent manner (Wang *et al*., 2016), fruits from strawberry plants sprayed with chitosan and MeJ on three occasions before harvesting showed increased postharvest resistance against *B. cinerea* infection (Saavedra *et al*., 2017), and strawberry plants sprayed with BTH showed induced resistance against *Phytophthora cactorum* and powdery mildew (*Podosphaera macularis*) (Eikemo, Stensvand and Tronsmo, 2003; Hukkanen *et al*., 2007).

All the reports on defence priming in strawberry cited above used plants that were either subjected to initial cold storage and/or were treated with pesticides to keep the plants disease-free. Thus, although these studies demonstrate primed or upregulated defences in strawberry they also confound any priming effect with the effect of subjecting the plants to pesticide treatment or cold storage. Cold treatment and pesticide application can affect plant physiology and may possibly have a priming-like effect. For example, cold treatment is known to induce epigenetic modifications and molecular memories that may be inheritable across generations (Yang, Howard and Dean, 2014; Berry and Dean, 2015; Dubin *et al*., 2015; Tao *et al*., 2017; Yang *et al*., 2017). The use of pesticides also defy the purpose of investigating induced plant defences, as defence priming agents should serve as an alternative to pesticide treatment.

Understanding the transcriptional reprogramming of *F. vesca* upon pathogen infection will identify key genes and pathways that help in devising appropriate agronomic solutions. In addition, exploiting the knowledge of pathogen induced transcriptional responses to choose appropriate defence priming chemicals will be a valuable method for crop protection schemes. Furthermore, systematic analysis of the effect of different priming chemicals with different treatment methods has not been investigated in strawberry. Analysis of the plants’ responses to the priming chemicals in the context of transcriptional reprogramming of *F. vesca*, defence responses and plant growth will provide substantial insights into the mode of action of these chemicals.

With this background, this study investigates: (1) transcriptional reprogramming of *F. vesca* leaves upon *B. cinerea* infection; (2) effect of four defence priming agents (BABA, MeJ, ProCa and RBH) to *B. cinerea* infection and plant growth; and (3) long-term effects on resistance of treating seeds with defence priming agents. Our results provide novel insights into the responses of *F. vesca* to *B. cinerea* infection and the effects of defence priming agents in providing protection against disease.

## 3. Results

### 3.1 *Botrytis cinerea* induces major transcriptional reprogramming in *Fragaria vesca*

To provide baseline information about transcripts involved in *F. vesca* defenses we studied the transcriptional reprogramming in *F. vesca* infected by *B. cinerea* using RNA-seq analysis of leaves from five infected and five mock-treated plants harvested 24 hours after infection. An average of 26,176,008 reads were generated per sample (median: 25,291,981) and the data were submitted to GenBank as XXX. Out of 28,588 mapped transcripts, 7003 were significantly differentially expressed in infected plants (p-value < 0.05) (Table S1), representing 24% of the *F. vesca* transcriptome. Among these differentially expressed genes (DEGs), 3314 were downregulated and 3689 were upregulated after infection, indicating a major transcriptional response upon *B. cinerea* infection. As expected, many upregulated DEGs were involved in defence responsive hormone biosynthetic pathways. Examples include genes encoding phenylalanine ammonia-lyase (*PAL*) involved in the SA pathway, 12-oxophytodienoate reductase (*12OR*) involved in the JA pathway, and 1-aminocyclopropane-1-carboxylate oxidase (*ACCox*) involved in ethylene (ET) biosynthesis (Table S1). In addition, many genes involved in the biosynthesis of defence metabolites such as terpenoids, flavonoids and major allergens were upregulated, suggesting that several defence systems were activated upon pathogen attack (Table S1). Interestingly, we also observed upregulation of the genes encoding gibberellin precursor ent-copalyl diphosphate synthase (*ECDS*) and the DELLA protein RGL1 that acts as a repressor of gibberellin signalling (Table S2). To confirm the differential gene expression observed in the RNA-seq data, qPCR analysis of selected genes was performed (Table 1). The qPCR gene expression results showed similar regulation as the RNA-seq data, thereby confirming the RNA-seq results (Figure 1).

**Table 1:** List of genes selected from the RNA-seq dataset for qPCR expression analysis. The genes are grouped according to the annotated functions, the gene IDs from the latest *F. vesca* 4.0 annotations and the corresponding log2-fold change values from the *B. cinerea* infected RNA-seq data set are provided.

| Gene groups | F_vesca_v4.0 | Gene Name | Description | log2FC | P adjusted |
| --- | --- | --- | --- | --- | --- |
| **Signaling and transcription factor** |  |  |  |  |  |
|  | FvH4_6g20130.1 | NHL10 | protein YLS9-like | 2.333 | 0.0001293 |
|  | FvH4_6g53770.1 | WRKY75 | probable WRKY transcription factor 75 | 4.957 | 4.402E-36 |
|  | FvH4_6g20840.1 | CPK1 | Transferring phosphorus-containing groups;<br>Calcium/calmodulin-dependent protein kinase | 3.292 | 7.763E-18 |
|  | FvH4_4g26910.1 | Lr10 | probable receptor-like protein kinase At1g67000 | 1.231 | 0.0011761 |
|  | FvH4_4g07370.1 | SLP3 | subtilisin-like protease SBT2.5 | -2.602 | 5.563E-10 |
|  | FvH4_4g35230.1 | ARR3 | two-component response regulator ARR5 | -2.919 | 1.516E-39 |
|  | FvH4_2g17220.1 | PHOT2 | putative LOV domain-containing protein | -2.049 | 1.668E-16 |
|  | FvH4_2g20290.1 | DSLPR | WD repeat-containing protein 70 |  |  |
| **Pathogenesis related** |  |  |  |  |  |
|  | FvH4_2g02920.1 | PR1 | pathogenesis-related protein 1-like | -1.374 | 0.0012965 |
|  | FvH4_3g05950.1 | PR4 | pathogenesis-related protein PR-4-like | 3.887 | 7.681E-15 |
|  | FvH4_6g16950.1 | PR5.3 | thaumatin-like protein | 2.195 | 3.298E-05 |
|  | FvH4_3g28390.1 | BG2-1 | glucan endo-1,3-beta-glucosidase-like | 3.092 | 6.34E-21 |
|  | FvH4_4g19500.1 | BG2-3 | glucan endo-1,3-beta-glucosidase, basic vacuolar isoform | 4.885 | 5.946E-39 |
|  | FvH4_6g22790.1 | PGIP1 | polygalacturonase-inhibiting protein | 1.728 | 7.187E-11 |
|  | FvH4_4g05760.1 | PecLy1 | probable pectate lyase 8 | -4.456 | 1.882E-26 |
| **Secondary metabolite and hormone pathway** |  |  |  |  |  |
| Salicylic acid | FvH4_7g19130.1 | PAL2 | phenylalanine ammonia-lyase | 2.539 | 4.549E-15 |
| Salicylic acid | FvH4_3g03130.1 | SCMT | salicylate carboxymethyltransferase-like | -2.784 | 3.206E-13 |
| Terpenoids | FvH4_6g11440.1 | GDS | (-)-germacrene D synthase-like | 6.279 | 3.137E-48 |
| Terpenoids | FvH4_6g36500.1 | BAS | beta-amyrin synthase | 3.485 | 8.188E-07 |
| Flavonoids | FvH4_6g16460.1 | 4CL7 | 4-coumarate--CoA ligase-like 7 | 6.767 | 3.431E-20 |
| Jasmonic acid | FvH4_5g32630.1 | 12OR2 | putative 12-oxophytodienoate reductase 11 | 2.773 | 1.942E-22 |
| Ethylene | FvH4_5g19290.1 | ACCox1 | 1-aminocyclopropane-1-carboxylate oxidase homolog 1-like | 2.881 | 1.032E-32 |
| Gibberellin | FvH4_2g23440.1 | ECDS | ent-copalyl diphosphate synthase, chloroplastic-like | 6.814 | 9.414E-55 |

**Figure 1:**
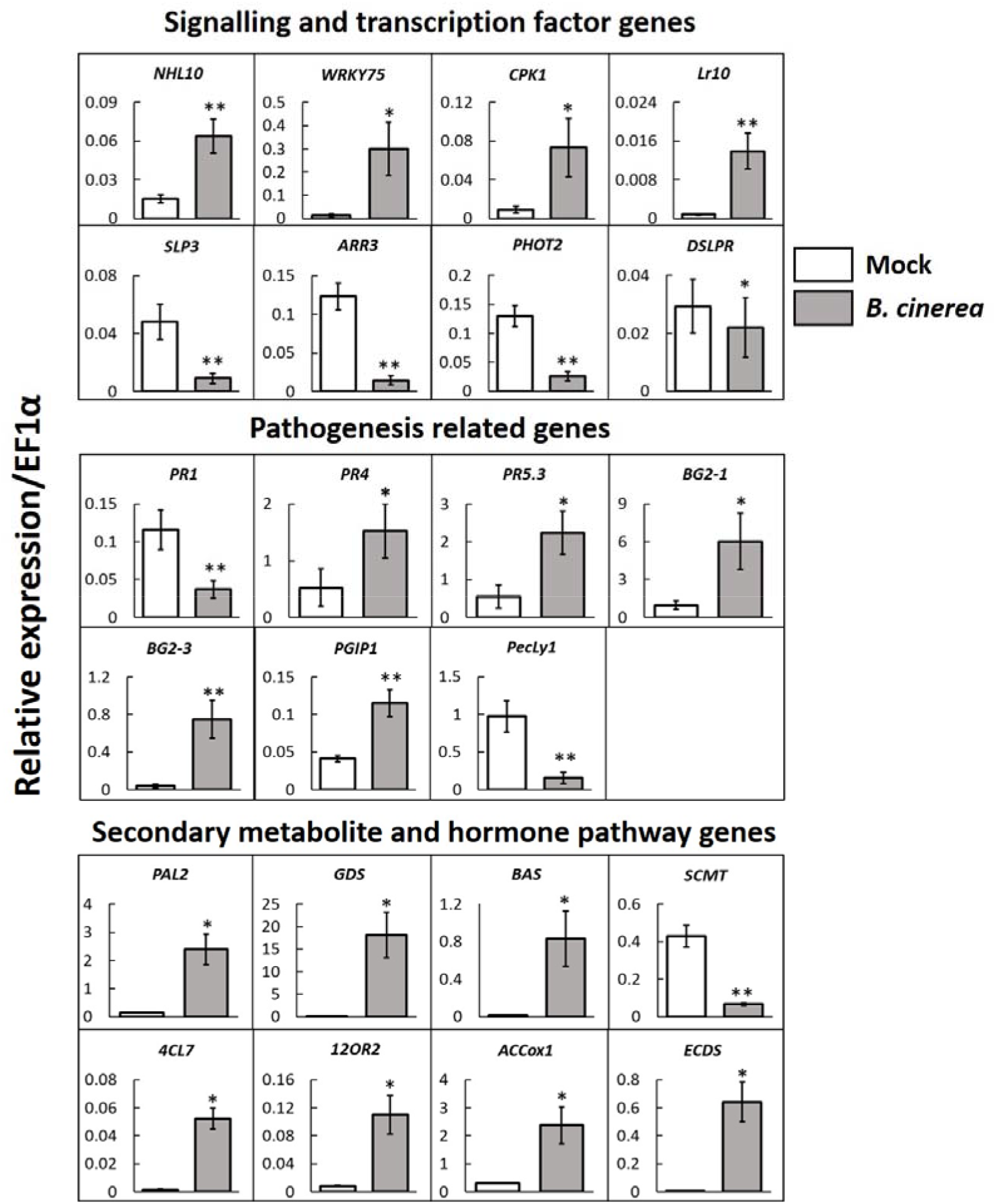
*Botrytis cinerea* induces transcriptional reprogramming in *Fragaria vesca*. Validation of expression levels of selected genes (Table 1) in *B. cinerea* infected and mock infected *F. vesca* leaves by qPCR. The housekeeping gene *EF1α* was used as an endogenous control gene and the expression values were calculated using the Δ-Ct method. *n=5*, student’s t-test: * *p < 0.05, **p < 0.01*.

#### 3.1.1 *Botrytis cinerea* and *Phytophthora cactorum* induce very similar transcriptomic responses in *Fragaria vesca*

To test if the transcriptomic response of *F. vesca* varies with the pathogen’s lifestyle and the plant organ that is infected we compared our RNA-seq data with previously published data from *F. vesca* roots infected with the oomycete hemibiotroph *Phytophthora cactorum* (Toljamo *et al*., 2016). In contrast, *B. cinerea* infects above-ground plant parts and is generally known to have a necrotrophic lifestyle, possibly with a short initial biotrophic phase (van Kan, Shaw and Grant-Downton, 2014; Veloso and Kan, 2018). Our comparison of DEGs in leaf and root tissues infected by *B. cinerea* (Table S1) or *P. cactorum* (Toljamo *et al*., 2016) respectively revealed great similarities in gene expression patterns. This prompted us to make a more detailed transcriptome comparison of strawberry’s responses to *B. cinerea* and *P. cactorum* infecting shoots and roots, respectively (Table S3). The overall similarity was visualised in a scatter plot comparing the transcriptional responses to the two pathogens. Expression patterns were positively correlated (r = 0.73; R^2^ = 0.54) for 4227 genes between *F. vesca* infected by *B. cinerea* (*Fv-Bc*) and *F. vesca* infected by *P. cactorum* (*Fv-Pc*), implying a high degree of similarity in the plant’s responses to the two pathogens (Figure 2a). A Venn diagram of up- and downregulated genes also illustrated the high number of genes that were similarly regulated between *Fv-Bc* and *Fv-Pc* (2251 + 1573 genes; Figure 2b). However, some genes displayed opposite regulation in the two systems (Figure 2a quadrant II and IV, Figure 2b), indicating differences in transcriptomic responses of different *F. vesca* organs to infection by different pathogens. Scatter plot analysis comparing transcriptomic responses of *Fv-Bc* and *F. vesca* infected by the ascomycete fungus *Colletotrichum acutatum* (*Fv-Ca*) (Amil-Ruiz *et al*., 2016) showed lower level of similarly regulated genes than between *Fv-Bc* and *Fv-Pc* (r = 0.24; R^2^ = 0.06) (Figure S1a and Table S4). Also, the transcriptomic responses of *Fv-Bc* and *A. thaliana* infected by *B. cinerea* (*At-Bc*) (Ferrari *et al*., 2007) showed a lower level of similarly regulated genes compared with *Fv-Bc* and *Fv-Pc* (r = 0.59; R^2^ = 0.34), probably owing to species differences (Figure S1b and Table S5).

**Figure 2:**
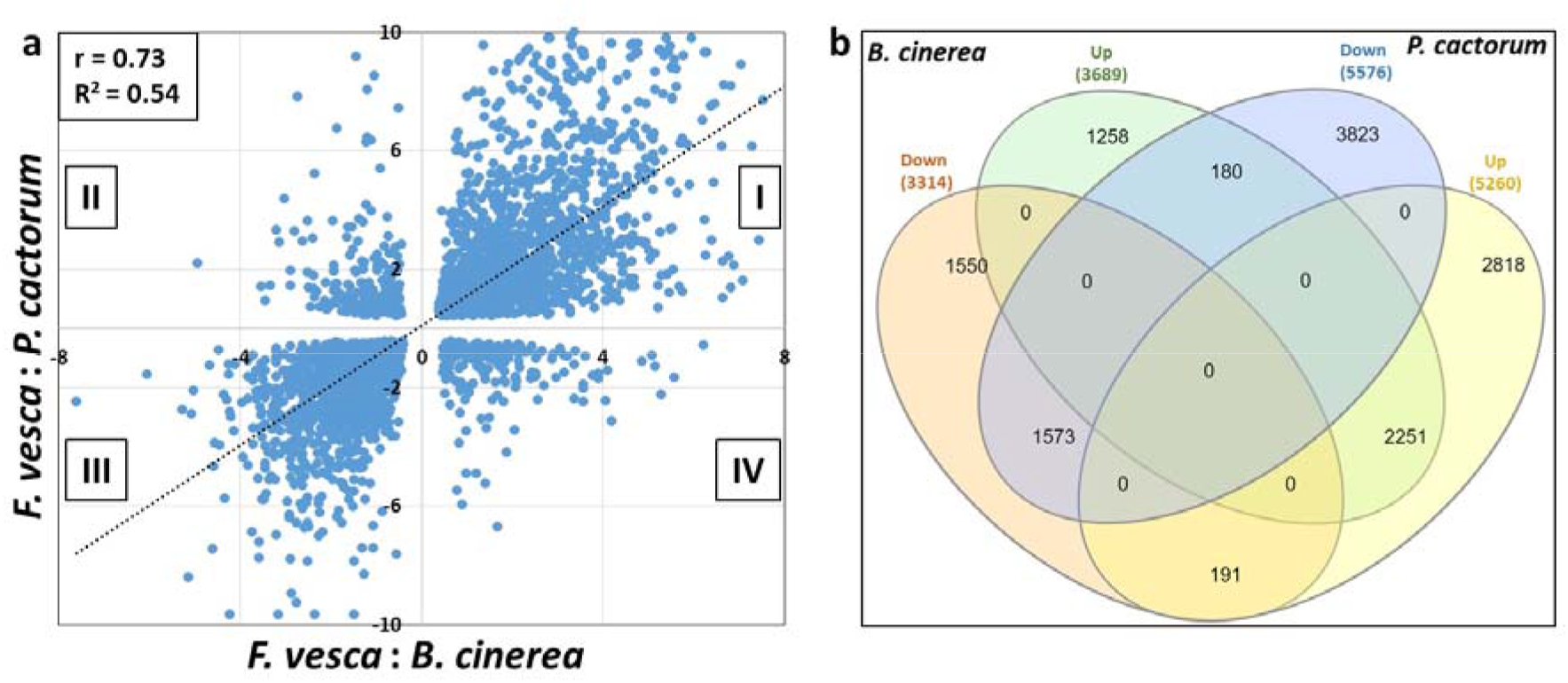
*Botrytis cinerea* and *Phytophthora cactorum* induce similar transcriptomic responses in *Fragaria vesca*. (a) Scatter plot of log_2_-fold change values of differentially expressed genes (DEGs) common between *B. cinerea* and *P. cactorum* infection. Each dot represents a single gene and shows the log_2_ FC-values of expression in strawberry plants infected with both *B. cinerea* and *P. cactorum*. The dots in quadrants I and III represent genes that respond similarly to both the pathogens, while the dots in quadrants II and IV represent the genes that respond oppositely to both pathogens. ‘r’ represents the correlation coefficient and ‘R^2^’ the coefficient of determination (b) Venn diagram showing numbers of DEGs common between *B. cinerea* and *P. cactorum* infection.

### 3.2 Priming agent-induced defence responses of *Fragaria vesca* against *Botrytis cinerea*

#### 3.2.1 Effects of defence-priming agents on *Fragaria vesca* resistance to *Botrytis cinerea*

The RNA-seq data showed upregulation of several defence related genes upon *B. cinerea* infection, including genes involved in SA, JA and ET signalling and synthesis of secondary metabolites (Table S2). As the priming agents BABA, MeJ and RBH normally function by inducing these defence responsive pathways, we investigated if BABA, MeJ and RBH also activated such defence pathways in *F. vesca* and increased resistance against *B. cinerea*. We treated plants with potential defence priming agents two days prior to fungal inoculation. Soil-drench treatment with RBH induced resistance in *F. vesca*, as the relative amounts of *B. cinerea* was less than half in the leaves of RBH treated plants compared to the control (Figure 3). However, BABA did not increase resistance, but on the contrary seemed to induce susceptibility or decrease resistance, as relative *B. cinerea* amounts were about twice as high in leaves of BABA treated plants as compared to the control (Figure 3). In Arabidopsis, BABA treatment has been found to confer protection against both *B. cinerea* (Zimmerli, Métraux and Mauch-Mani, 2001) and abiotic stresses like salt and drought (Jakab *et al*., 2005). To verify our observation with BABA-soil drench, we tested different concentrations of BABA and observed that plants had increasing amounts of *B. cinerea* infection with increasing BABA concentrations (Figure S3). BABA treatment combined with drought stress reduced the relative water content (RWC) of *F. vesca* in a BABA-concentration dependent manner (Figure S4), indicating that BABA increased susceptibility to both types of stress, biotic and abiotic.

**Figure 3:**
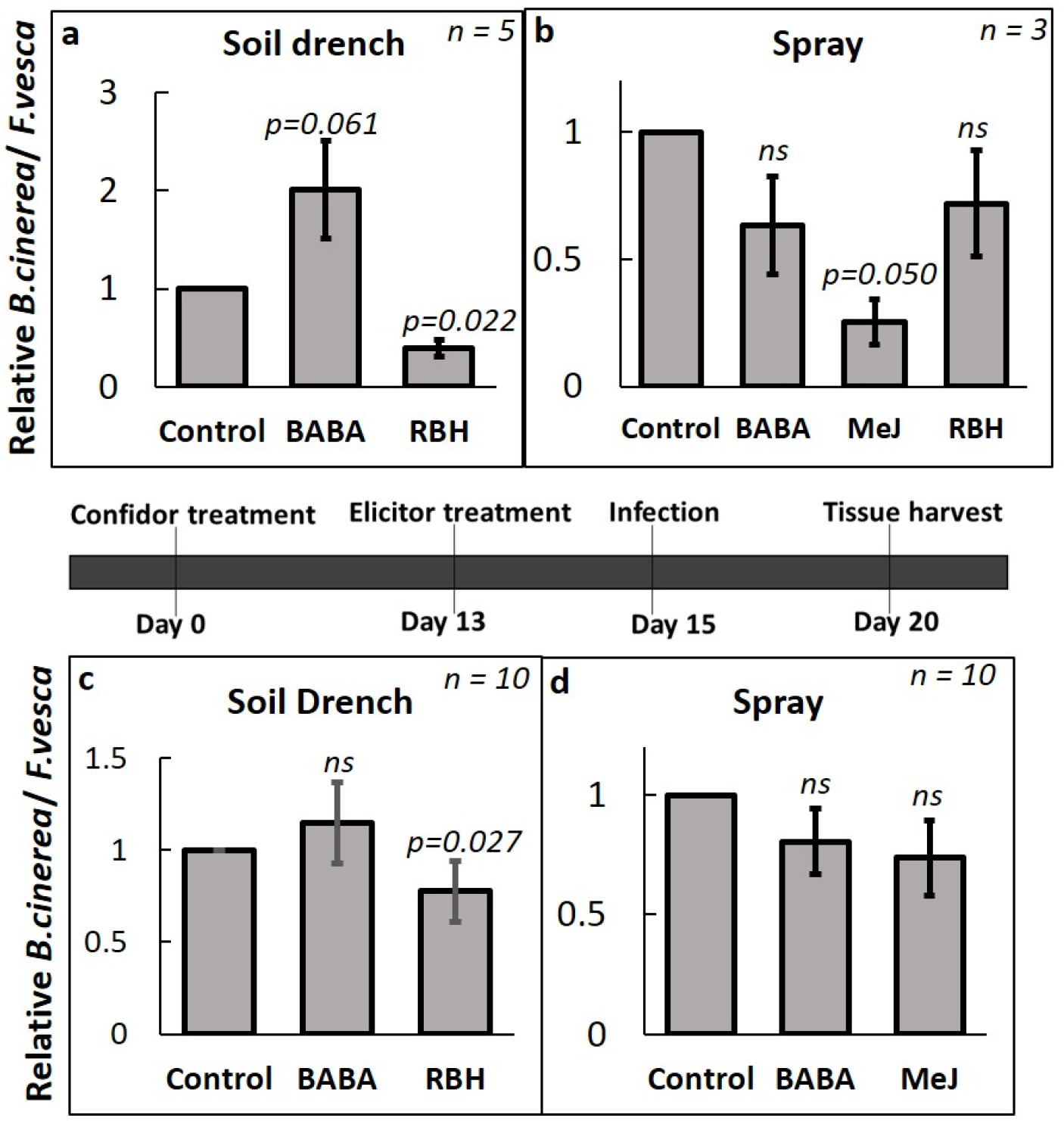
Effects of priming chemicals on *Fragaria vesca* resistance against *Botrytis cinerea*. (a) Amount of *Botrytis cinerea* quantified by qPCR after treatment with priming agents by soil-drenching (b) spraying. (c-d) plants treated with Imidacloprid before priming agent treatments. ‘*n*’ represents number of biological replicates. Error bars indicate standard error. Student’s t-test was performed for statistical significance. ‘*ns*’ represents not statistically significant.

Spray treatment of BABA, MeJ and RBH all decreased *B. cinerea* infection but only MeJ treatment was statistically significant, indicating that these chemicals primed or induced resistance in *F. vesca* (Figure 3b). Extensive growth of *B. cinerea* on potato dextrose agar plates with high concentrations of these chemicals suggests that the results *in planta* were not due to direct effects of the chemicals on *B. cinerea* growth and survival (Figure S5).

To test the effect of putative defence priming agents under the influence of a plant protection chemical, *F. vesca* plants were treated with an insecticide Imidacloprid (Confidor) two weeks before treatment with priming agent. As shown in Figure 3c-d, *B. cinerea* amounts differed little between treated plants and control plants, both for soil drench and foliar spray (Figure 3a-b). For example, plants treated with Imidacloprid before BABA were not significantly different from the control plants.

Plant growth was significantly inhibited by soil drench treatment with BABA and RBH in non-infected plants (Figure 4a). Spray treatment with BABA, MeJ and RBH did not have any significant effects on the growth rate of plants (Figure 4b).

**Figure 4:**
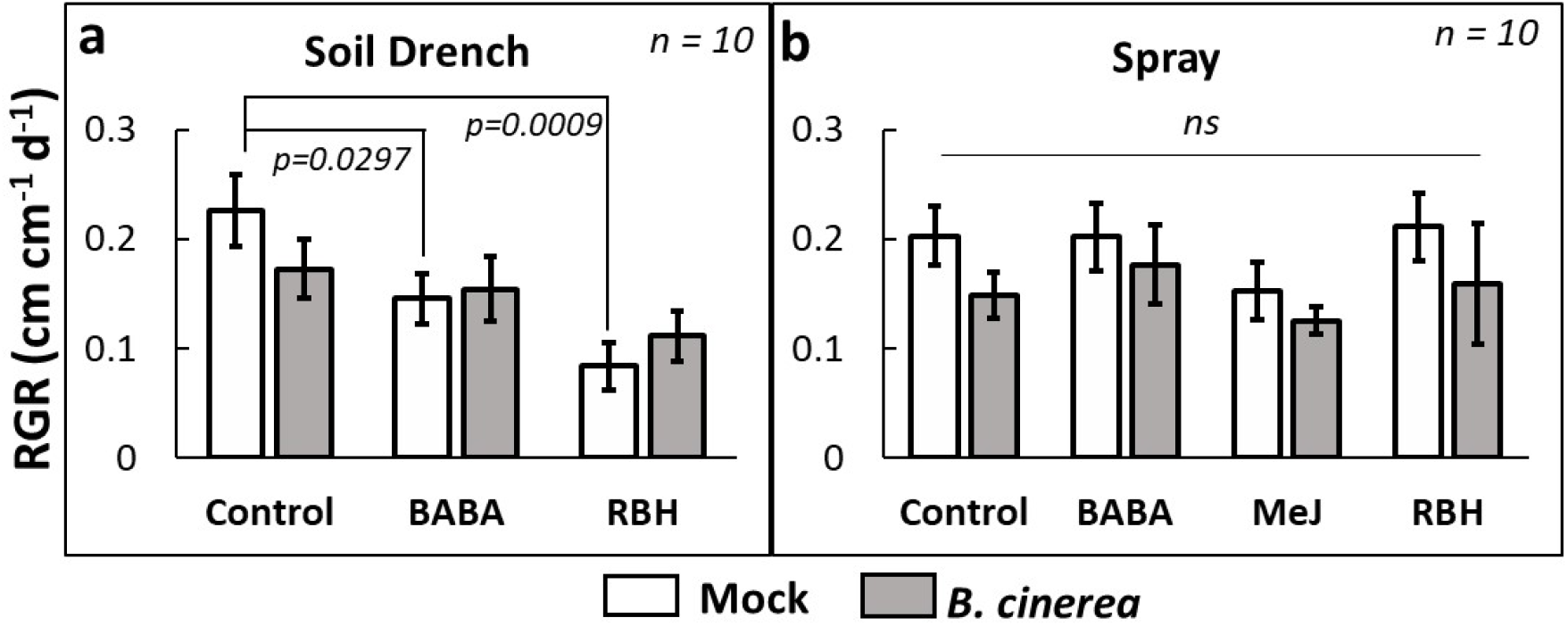
Effect of priming chemicals on relative growth rate (RGR) of *Fragaria vesca*. RGR of soil-drenched (a) and spray treated (b) plants subsequently infected with Botrytis spores or mock control. ‘*n*’ represents number of biological replicates. Error bars indicate standard error. Student’s t-test was used for statistical significance. ‘*ns*’ represents not statistically significant.

#### 3.2.2 Gibberellic acid and ProCa influence resistance against *Botrytis cinerea*

The RNA-seq data showed that the gibberellin precursor encoding gene *ECDS* was upregulated upon *B. cinerea* infection. We investigated if ProCa induces resistance against *B. cinerea* infection in *F. vesca. Fragaria vesca* plants that were sprayed twice with prohexadione-calcium (ProCa) before inoculation showed more visible necrosis than mock treated control plants (Figure S6). Quantification of *B. cinerea* levels in infected leaves showed that ProCa-treated plants had 12-fold higher *B. cinerea* levels than the control (Figure 5a), indicating reduced resistance. To investigate if ProCa and GA have antagonistic effects on resistance against *B. cinerea*, *F. vesca* plants were sprayed once with either ProCa (100 mg/L) or GA_3_ (50 mg/L) before inoculation. Plants sprayed with GA_3_ were more resistant to *B. cinerea* than the controls, whereas plants sprayed with ProCa were less resistant (Figure 5b). GA_3_ also increased the growth of mock treated control plants, while ProCa had little effect on growth (Figure 5c). These results indicate that GA levels in *F. vesca* are important for resistance against *B. cinerea*.

**Figure 5:**
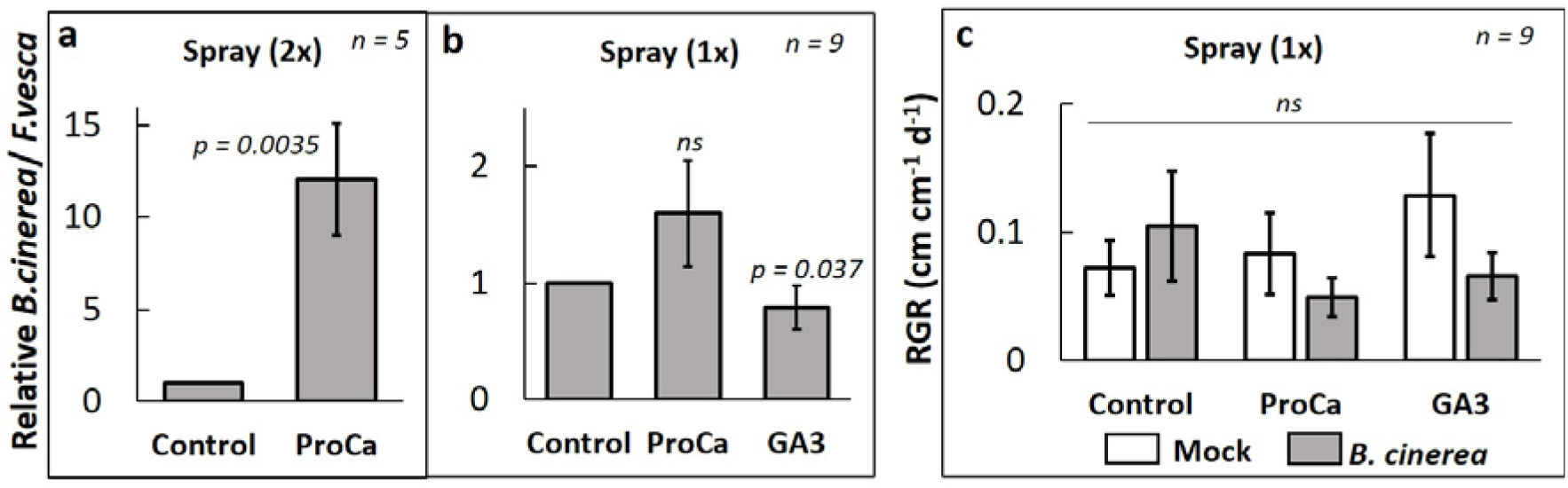
Effect of gibberellin biosynthesis modulators on resistance against *Botrytis cinerea*. (a) Relative *B. cinerea* amounts in two times (2x) spray treated Prohexadione-calcium (ProCa) (100 mg/L). (b) Relative *B. cinerea* amounts in plants one time (1x) spray treated with ProCa and Gibberellic Acid (GA_3_) (50 mg/L). (c) Relative Growth Rate (RGR) analysis of the plants spray treated with either ProCa or GA_3_ and with or without *B. cinerea* inoculation. ‘*n*’ represents number of biological replicates. Student’s t-test was used for statistical significance. ‘*ns*’ represents not statistically significant.

#### 3.2.3 Seed priming for induced resistance against *Botrytis cinerea*

We then studied the effect of seed priming on germination, vegetative growth and resistance against *B. cinerea*. Seeds of *F. vesca* treated overnight with BABA, MeJ and RBH did not show differences in germination rates as compared to control treatment (Figure 6a). While treatment with BABA and MeJ had no significant effect on early growth of seedlings compared with control, RBH-treated seeds showed higher petiole lengths, indicating a positive effect of RBH on early seedling growth (Figure 6b). Plants grown from seeds treated with BABA and MeJ were more susceptible to *B. cinerea* infection than control treatment whereas plants from RBH treated seeds were more resistant to infection, indicating induced or primed-resistance following seed treatments with RBH (Figure 6c). Seedlings from MeJ-treated seeds showed some temporary growth inhibition 12 days after germination (Figure S7), but this effect was not significant in the longer term (Figure 6b, d). Interestingly, RBH seed treated and infected seedlings grew significantly better than mock-inoculated seedlings during the six days of infection period (Figure 6d).

**Figure 6:**
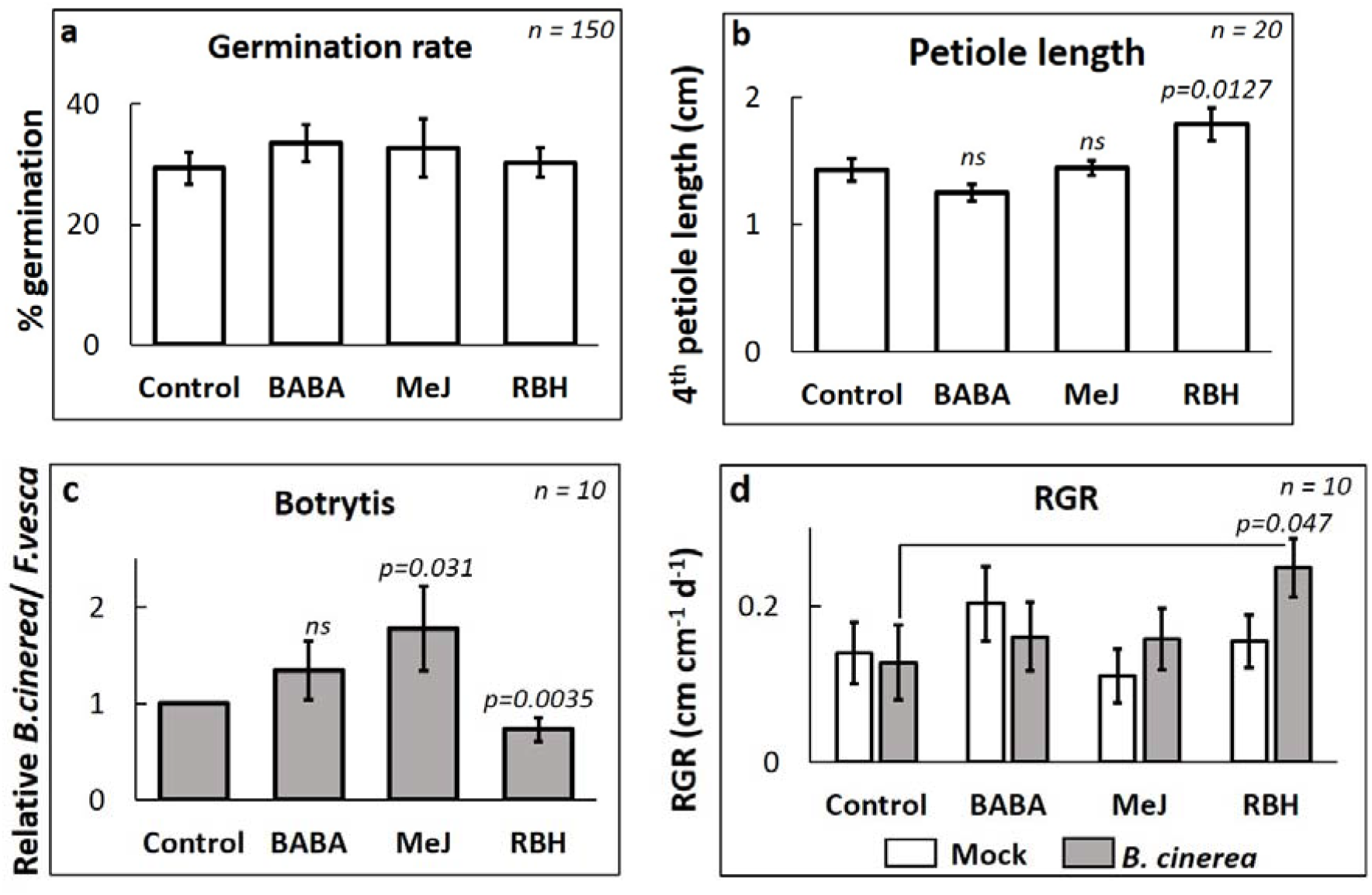
Effect of seed treatment of *Fragaria vesca* on germination, vegetative growth and resistance against *Botrytis cinerea*. (a) Germination rates of seeds treated with either β-aminobutyric acid (BABA), methyl jasmonate (MeJ) or (*R*)-β-homoserine (RBH). (b) The length of 4th petioles of 5-week old seedlings from seeds treated with BABA, MeJ and RBH. (c) Relative *B. cinerea* amounts in plants grown from priming agent treated seeds and infected with *B. cinerea*. (d) Relative Growth Rate (RGR) analysis of young petioles of plants grown from seeds treated with BABA, MeJ and RBH. ‘*n*’ represents number of biological replicates. Error bars indicate standard error. Statistical significance was tested using student’s t-test. ‘*ns*’ represents not statistically significant.

#### 3.2.4 RBH primes specific defence genes

To understand the molecular details of the observed priming effect of RBH seed treatments, gene expression analysis of the four chosen genes were performed with and without infection. The genes PR4, *BG2-3, 4CL, ECDS*, from different pathways were chosen from our RNA-seq data and show upregulation after infection. As shown in Figure 7, *PR4* is only slightly upregulated upon RBH treatment, but highly upregulated after RBH treatment and *B. cinerea* infection, a signature of priming. Seed treatments with BABA did not have effects on *PR4* and *BG2-3* but *4CL* and *ECDS* were slightly upregulated upon BABA treatment alone and induced upon infection. These observations indicate that priming by RBH and BABA have some overlaps and some differences that is important for revealing resistance mechanisms against *B. cinerea*. The resistance induced by RBH alone suggests that the specific genes/pathways primed by RBH are important for resistance against *B. cinerea*.

**Figure 7:**
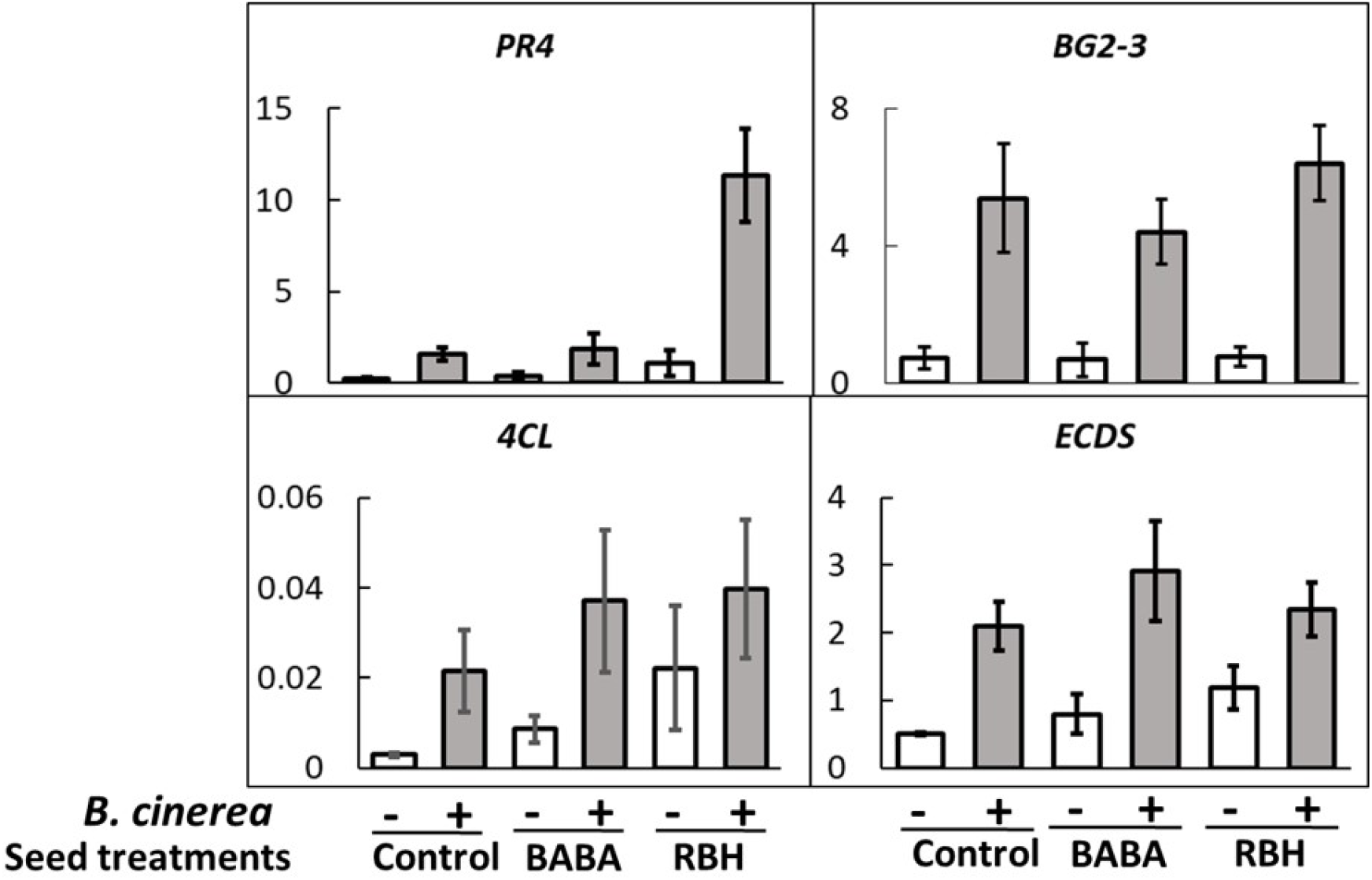
Gene expression analysis by qPCR of selected genes from Table 1 upon seed treatment and subsequent infection. The gene *PR4* is upregulated upon (*R*)-β-homoserine (RBH) treatment only and shows boosted expression upon Botrytis infection, at least 6 fold higher than the control and β-aminobutyric acid (BABA) treatments. The genes *BG2-3, 4CL* and *ECDS* show similar upregulation upon infection in all the treatments, *n=3*.

## 4. Discussion

To effectively address problems related to plant diseases it is important to understand how plants respond to pathogens. *Botrytis cinerea* is an important plant pathogen with a very broad host range. We found that *B. cinerea* induces large transcriptomic responses in *F. vesca* that include genes functioning in major hormone signalling and secondary metabolite pathways (Table S1). Interestingly, we observed many similarities in the transcriptomic responses of *F. vesca* leaves infected by *B. cinerea* (*Fv-Bc*) and *F. vesca* roots infected by *P. cactorum* (*Fv-Pc*) (Toljamo *et al*., 2016). Thus, the recognition mechanisms, downstream signalling and genes that *F. vesca* uses to resist infection by these two pathogens appear to be similar. In Figure 2, quadrant I and III show genes that are similarly regulated in the *Fv-Bc* and *Fv-Pc* system and quadrant II and IV show genes that are oppositely regulated. It is interesting to see that two different pathogens infecting above-ground and below-ground tissues induce similar transcriptional reprogramming. Comparison of *F. vesca* leaves infected by *B. cinerea* or *C. acutatum* did not show such similarities (Amil-Ruiz *et al*., 2016; Figure S1), despite the fact that both *B. cinerea* and *C. acutatum* are necrotrophic leaf pathogens.

These observations indicate that similarities and differences in the pathogen perception and the downstream plant responses might be independent of the pathogen’s lifestyle. Whether similar plant’s responses mean similar effectiveness of the crop protection measures is an interesting question that needs investigation. For example, it can be envisaged that measures for protection against *B. cinerea* might also be effective against *P. cactorum* because the *F. vesca* plant perceives and responds similarly to both the pathogens. However, experiments to uncover these responses are required to confirm these speculations and to develop effective crop protection schemes. Comparison of *Fv-Bc* with *At-Bc* (Arabidopsis infected with *B. cinerea*) revealed interesting insights (Figure S1b). Although, there are similarities in the regulation of homologous genes, we observed some interesting differences. One prominent difference was the expression of a key resistance gene, *Pathogenesis-Related 1* (*PR1*), that is highly induced upon *B. cinerea* (Ferrari *et al*., 2007) and *Pseudomonas syringae* pv *tomato* DC 3000 (*Pst* DC 3000) infection in Arabidopsis (Zimmerli *et al*., 2000), whereas the corresponding *F. vesca* ortholog (*FvPR1.1*) is downregulated upon infection by *B. cinerea* (Table S2) and *P. cactorum* (Toljamo *et al*., 2016). Whether these differences stem from the plant’s own responses or if the pathogen manipulates the host’s immune responses differently is also an interesting question that needs further investigations.

Genes involved in well-known defence responsive pathways such as jasmonic acid (JA), salicylic acid (SA), ethylene (ET) and terpenes were upregulated upon *B. cinerea* infection. Defence priming agents such as BABA, MeJ and RBH impart resistance by inducing the defence responsive pathways such as SA, JA and ET. Primarily using BABA and RBH, we explored the effects of different treatment methods in imparting resistance against *B. cinerea*. Soil-drench treatment of BABA induces susceptibility and spray treatment of the BABA induces resistance against *B. cinerea* (Figure 3a-d and Figure S3). As the infection assays are performed in leaves, it follows that BABA induces resistance to the treated plant part and not the parts distant to the treatments, reflecting the local resistance rather than systemic resistance. Drought experiments using BABA soil-drenched plants indicate that BABA decreases the ability to retain water in the leaves, also following similarities in responses as infection by *B. cinerea*. These observations were consistent between *F. vesca* Hawaii-4 and *F. vesca* Alexandria ruling out the genotype specific differences. Whereas for RBH, both soil-drench and spray treatments imparted resistance, with soil-drench being most effective, probably reflecting that systemic resistance is more effective than local resistance. RBH imparts resistance by cell-wall reinforcement and inducing JA and ET pathways in Arabidopsis (Buswell *et al*., 2018). However, if the mechanisms of RBH-induced resistance (RBH-IR) in *F. vesca* are similar to that in Arabidopsis needs further investigations. Understanding if RBH-IR is active for several weeks after soil-drench or spray treatment will hold great potential for agricultural applications.

A report in Arabidopsis (Zimmerli *et al*., 2000) indicates the formation of necroses after spray treatment of BABA, that can induce systemic acquired resistance (SAR) pathway. Such induction of SAR pathway can mask the primary effect of BABA, forming the reason for not using the spray treatment in Arabidopsis (Zimmerli *et al*., 2000). However, no visible necroses was observed in *F. vesca* plants upon spray treatment of chemicals or the mock treatment. Also, there are reports of spray treating strawberry plants with different chemicals to study induced resistance (Terry and Joyce, 2000; Eikemo, Stensvand and Tronsmo, 2003; Hukkanen *et al*., 2007; Landi *et al*., 2017; Saavedra *et al*., 2017). We therefore consider the observed effects to be the plant’s induced responses to priming agent treatments.

The experiment with an insecticide imidacloprid showed similar effects of BABA and RBH as in the experiments with untreated plants, however with reduced effectiveness, possibly indicating the interference of imidacloprid with the functions of priming chemicals. Based on this observation, we argue that using chemical control agents for growth and maintenance of plants interferes with the studied plant’s responses and may provide less reliable results, especially when studying effectiveness of defence priming agents. These observations therefore serve as a cautionary note to the researchers studying the effects of priming agents and emphasizes the importance of using clean, healthy and non-chemical treated plants in studying plant responses to defence priming agents.

Our experiments with ProCa provided some interesting insights. ProCa acts by blocking the dioxygenases that require 2-oxoglutaric acid as a co-substrate in gibberellin biosynthesis. Due to this action, the plants treated with ProCa accumulate less bioactive gibberellins that are required for shoot growth, thereby display shoot growth inhibition (Rademacher, 2004; McGrath *et al*., 2009). Spray treating plants twice before *B. cinerea* inoculation increased the infection spread in contrast to the reported observations against necrotrophic pathogens in the literature. ProCa’s ability to induce resistance in Rosaceae plants against necrotrophs makes our observations more intriguing as *F. vesca* also belongs to the Rosaceae family. These results suggest that, in *F. vesca*, ProCa is probably not inducing the flavonoid antimicrobial compounds like in other Rosaceae plants. However, this only explains the absence of induced resistance and not the observed induced susceptibility. Reports in *F. vesca* suggest that application of ProCa inhibits runner formation and that subsequent GA_3_ application reverses this effect (Hytönen *et al*., 2009), pointing out the antagonistic functions of ProCa and GA_3_ when applied exogenously. Following these studies, our observations indicate that spray treatment of ProCa decreases while GA3 increases resistance against *B. cinerea* (Figure 5b).

The role of gibberellin pathway in defence responses is identified in Arabidopsis. Gibberellin triggers the degradation of DELLA proteins, a group of plant growth repressors. In Arabidopsis, it is observed that a quadruple knockout mutant of DELLA genes (quadruple-DELLA mutant), lacking four out of the five DELLAs, is resistant to the hemibiotroph *Pseudomonas syringae* pv. tomato strain DC3000 (*Pto* DC3000) and susceptible to the necrotrophic fungus *Alternaria brassicicola*. This is because the quadruple-DELLA mutant displayed attenuated induction of a JA responsive gene *PDF1.2* that is essential for JA mediated resistance against necrotrophs (Navarro *et al*., 2008). In rice, a necrotrophic fungus *Gibberella fujikuroi* (*Fusarium moniliforme*) causes *bakanae* or foolish-seedling disease leading to elongated seedlings and slender leaves causing drastic reduction in crop yields (YABUTA and T., 1938). Although it is known that GA produced by *G. fujikuroi* is responsible for the observed symptoms in rice, the mechanism of its virulence action is unknown. The authors (Navarro *et al*., 2008) propose that *G. fujikuroi* triggers the degradation of DELLAs by secreting GA into the plant, thereby disabling JA-mediated resistance against necrotrophs. On the contrary, exogenous application of GA decreased resistance against the hemibiotrophs, *Magnaporthe oryzae* (*Mo*) and *Xanthomonas oryzae* pv. oryzae (*Xoo*) in rice (Yang *et al*., 2008; Qin *et al*., 2013; De Bruyne, Höfte and De Vleesschauwer, 2014).

Our experiments indicate that spray application of GA_3_ induces resistance in *F. vesca* to the necrotrophic fungus *B. cinerea*, contrary to the observations in Arabidopsis and rice. In addition, *B. cinerea* infection induces upregulation of a gene encoding ent-copalyl diphosphate synthase (*ECDS*, ent-kaurene synthase A) involved in the biosynthesis of diterpenoid compounds and gibberellin (Table S1). Interestingly, *ECDS* is also upregulated in *F. vesca* upon infection by a hemibiotrophic oomycete *Phytophthora cactorum* (Toljamo *et al*., 2016), pointing out to the similarities in infection strategies and/or *F. vesca*’s responses between two different pathogens in the same host.

Treatment of *F. vesca* seeds with the priming chemicals BABA, MeJ or RBH did not affect seed germination and growth of the seedlings (Figure 6a-c), however only RBH treatment induced resistance against *B. cinerea* infection (Figure 6d). Interestingly, induced resistance against *B. cinerea* was partially consistent with that reported for tomato seed treatment. Tomato plants from seeds treated with JA are resistant against *B. cinerea* whereas plants from seeds treated with BABA are susceptible (Worrall *et al*., 2012). Also, consistent with the tomato seed treatment study (Worrall *et al*., 2012), we observed inhibition of seedling growth 12 days after germination upon MeJ treatment (Figure S6), which was not significant in the long term (Figure 6b, d).

The observation that seed treatments with RBH improves the ability of *F. vesca* to resist *B. cinerea* infection indicates that a priming signal is transmitted from the seed to the adult plant, reflecting molecular memory. Gene expression experiments (Figure 7) reveal that *BG2-3, 4CL* and *ECDS* are induced by both BABA and RBH seed treatments whereas *PR4* is only induced upon RBH treatment, clearly differentiating between the mechanisms of two different priming agents. Based on our results (Figure 6c), it is reasonable to assume that BABA and MeJ might also induce molecular memory that is functionally different from that of RBH induced memory in imparting resistance against *B. cinerea*. The difference in functionalities of BABA and RBH are also true for the soil-drench experiments, where we observed induced susceptibility with BABA and induced resistance with RBH (Figure 3a and Figure S3). As soil-drench BABA treatment induces broad-spectrum resistance against pathogens with different lifestyles in Arabidopsis (Zimmerli *et al*., 2000; Zimmerli *et al*., 2001), our results indicate that this does not necessarily imply its similar functions in other plants. These observations might also indicate the differences in underlying defence responses and the BABA perception and signalling mechanisms of *F. vesca*. Our results in *F. vesca* serves as an example and underline the importance of characterizing the defence responses in crop plants in order to develop effective crop protection strategies.

Overall, in addition to providing interesting insights, our results raise new questions about the *F. vesca-B. cinerea* interaction. It is interesting and intriguing to envisage what determines BABA and GA to function differently even though *B. cinerea* infection induces the widely described defence responsive pathways. Perhaps, further work into quantifying GA and the defence metabolites (JA, SA and ET) is required to understand the transcriptomic responses of *F. vesca* in the context of induced resistance. Such insights might also reveal important determinants governing the defensive state of *F. vesca*. Furthermore, RBH has emerged as an effective chemical agent in inducing resistance against *B. cinerea*, independent of treatment methods (soil-drench, spray and seed treatments), indicating its high potential for use as a priming agent for strawberry. Further research to test if RBH-priming is effective against broad spectrum of stresses in *F. vesca* will be valuable knowledge that will be of direct benefit to the agriculture sector.

## 5. Experimental procedures

### 5.1 Plant growth

*Fragaria vesca* ‘Hawaii-4’ and *F. vesca* ‘Alexandria’ accessions were cultivated in a growth room with a 14 h day (∼100 μmol m^−2^ s^−1^ photosynthetically active radiation (PAR) at 24 °C) and 10 h night (19 °C) at 40-45% relative humidity. Plants were grown in topsoil in 400 mL pots and were not subjected to any kind of chemical treatments (pesticides, fungicides or fertilizers) to ensure that the phenotypic responses we observed were caused by the defence priming chemicals alone. For priming and infection experiments, 8- to 12-week-old plants grown from seeds were used to ensure synchronous growth within each experimental batch. The plants were about 10-12 cm tall from the soil surface at the start of the experiments, bearing 6-10 trifoliate leaves. For the imidacloprid (Confidor) experiment, *F. vesca* ‘Hawaii-4’ seeds were germinated in greenhouse conditions and imidacloprid was administered to the plants through water two weeks before application of the priming agent.

### 5.2 Treatment with defence priming chemicals

Solutions of defence priming chemicals were prepared as follows: β-aminobutyric acid (BABA, Sigma-Aldrich, #A44207-5G), R-β-homoserine (RBH, Aurum Pharmatech, #S-4676), and prohexadione-calcium (ProCa, Sigma, #31720) were dissolved in distilled H_2_O, methyl jasmonate (MeJ, Sigma-Aldrich, #392707) was prepared in 0.1% Tween-20, and gibberellic acid (GA3, Duchefa, #G0907.0001) was first dissolved in 50 μl ethanol and then diluted in H_2_O. Plants were treated in three different ways: (i) soil-drench using ten times concentrated (10×) solutions of the priming chemical (Jakab *et al*., 2005; Wilkinson *et al*., 2018), (ii) spray treatment of above-ground parts using final concentration (1×) chemical solutions (Saavedra *et al*., 2017), and (iii) treatment of seeds. For soil-drench treatments, 10× solutions were prepared corresponding to the concentrations given in the figure legends. Plants were treated by pouring 10% of the pot volume of the 10× solution into each pot. The same volume of dH_2_O or 0.1% Tween-20 solutions were used for control treatments. For foliar sprays, solutions corresponding to the concentrations given in the figure legends were sprayed onto the plants using a hand-sprayer until run-off. Control plants were sprayed with dH_2_O or 0.1% Tween-20. Experiments with ProCa were performed by spraying *F. vesca* plants with 100 mg/L solution. Similar or higher ProCa concentrations have been used to induce resistance in plants in several previous studies (Yoder, Miller and Byers, 1999; McGrath *et al*., 2009; Spinelli *et al*., 2010). For seed treatments, seeds were surface sterilized and incubated overnight with rotation in 0.3 mM BABA, 0.3 mM MeJ or 0.5 mM RBH. Seeds were then washed twice with water and germinated on Petri dishes with moist filter paper. The germinated seedlings were transferred to pots 12 days after seed treatment.

### 5.3 *Botrytis cinerea* inoculum preparation and infection

*Botrytis cinerea* isolate Bc101 was grown on potato dextrose agar plates in darkness. Spores on four- to five-week-old plates were harvested using liquid potato dextrose broth (PDB) media. The spore suspension was passed through a sterile 70 μM nylon mesh and centrifuged at 3000g for 1 min, before the spore pellet was re-suspended in fresh PDB media. Spores were counted using a haemocytometer and the suspension was adjusted to a concentration of 10^6^ spores mL^−1^. The spore suspension was supplemented with 0.02% Tween-20 and sprayed on plants with a hand sprayer. PDB media supplemented with 0.02% Tween-20 was used for mock inoculations. Sprayed plants were placed in a tray, covered with transparent polypropylene bags to maintain high humidity, and incubated in a plant growth room until symptoms appeared. Necrotic lesions on leaves usually started appearing 4-5 days after spore spraying. When clear symptoms were observed, all the leaves were harvested for DNA extraction and qPCR quantification of *B. cinerea* colonization.

### 5.4 Genomic DNA based quantification method for *Botrytis cinerea* infected leaves

To establish a working quantification method for *B. cinerea* infection of *F. vesca*, leaves were drop inoculated with *B. cinerea* spore suspension and incubated under high humidity to promote fungal infection. We detected clear symptoms of infection, in the form of necrotic lesions on leaves, 5 days after infection (Figure S2a). As expected, *B. cinerea* quantification using genome-specific primers for *B. cinerea* (Bc3.1F and Bc3.1R) and *F. vesca* (EF1αF and EF1αR) revealed that infected leaves contained much more *B. cinerea* than mock treated controls, indicating fungal disease progression in the necrotic lesions (Figure S2b).

### 5.5 Drought treatments

BABA-treated *F. vesca* plants were water deprived by withholding water from the day of BABA treatment. Leaves were harvested over a 10-day interval (0D, 3D, 6D, 8D and 10D) after BABA treatment to determine their relative water content (RWC). RWC was calculated using the formula RWC = (FW – DW) × 100/(SW – DW) (So *et al*., 2014), where FW = fresh weight, SW = water saturated weight, and DW = dry weight of the leaves.

### 5.6 Gene expression and *Botrytis cinerea* quantifications

For each plant, about 100 mg of leaf tissue was ground in liquid N_2_ and used for DNA or RNA extraction. For DNA extraction, the DNeasy Plant Kit (#69106) was used in a QIAcube automated sample prep system according to the manufacturer’s instructions. Total RNA was isolated using Spectrum™ Plant Total RNA Kit with minor modifications (Badmi *et al*., 2018). Briefly, 100 mg of tissue powder was mixed with preheated lysis buffer containing CTAB (2%), PVPP (2%), Tris-Cl (pH 8.0, 100 mM), EDTA (pH 8.0, 25 mM), NaCl (1 M) and β-mercaptoethanol (1%). The mixture was incubated at 65 °C for 8 min, with vortexing for the first 60 seconds. The lysate was then centrifuged for 10 min at 13000 rpm, the supernatant was mixed with an equal volume of chloroform: isoamyl alcohol (24:1) and centrifuged for 10 min at 4 °C. The supernatant was transferred to the kit’s filtration column (blue retainer ring), and from this step we followed the manufacturer’s instructions. On-column DNaseI treatment was performed to ensure DNA-free total RNA. Total RNA (500 ng) was used to prepare cDNA using iScript™cDNA Synthesis Kit (#170-8891). SsoAdvanced™ Universal SYBR^®^ Green Supermix (#1725271) was mixed with genomic DNA or cDNA to perform qPCR on the Applied Biosystems ViiA 7 system to determine the C_T_ values for each primer pair.

Quantification of *B. cinerea* was performed using genomic DNA isolated from infected strawberry tissues. C_T_ values from *B. cinerea* specific genomic DNA primers (Bc3F and Bc3R, Table 2) were normalised against *F. vesca* specific *EF1α* primers to obtain the relative *B. cinerea* levels in the tissue. For RNA-seq validation using qPCR, *EF1α* was used as the housekeeping control gene and relative expression levels of genes were determined using the ΔC_T_ method (Pfaffl, 2001).

**Table 2:**
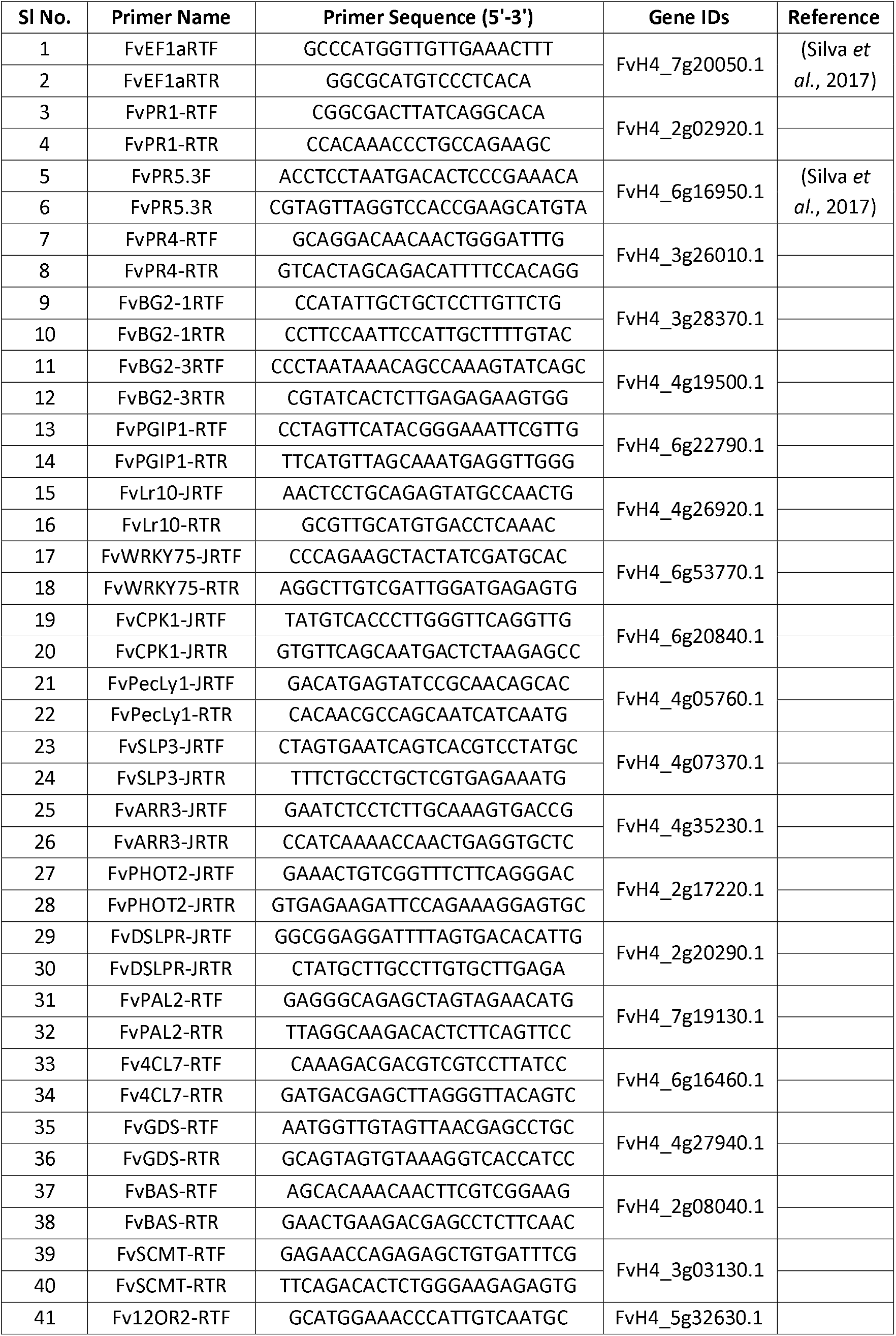
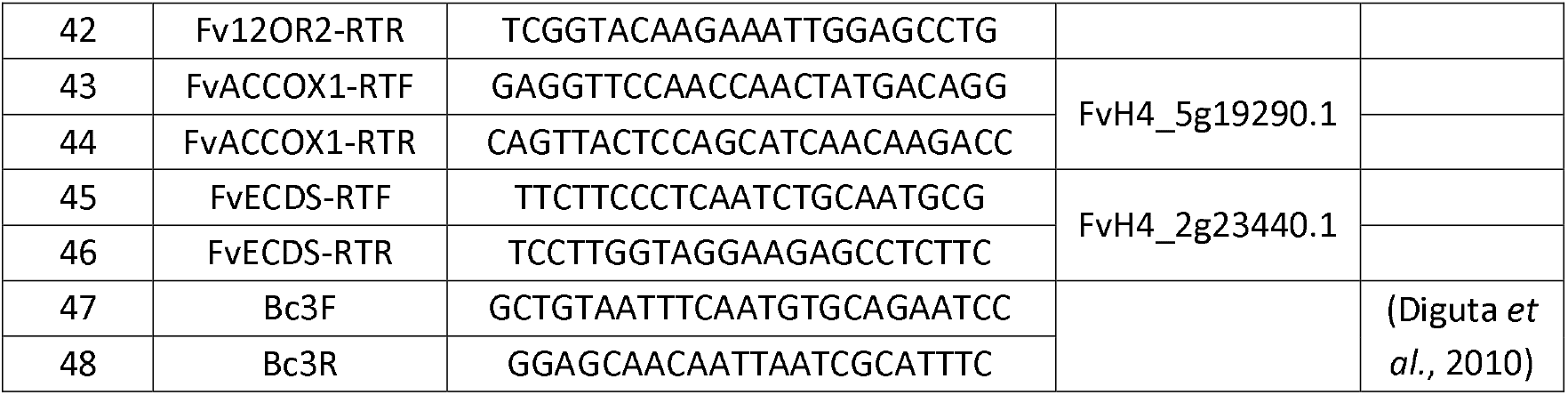
List of primers used in this study.

### 5.7 RNA-seq analysis

To study the transcriptomic responses of *F. vesca* to *B. cinerea* infection, plants were infected with *B. cinerea* spores as described above and harvested 24 hours later for RNA isolation. Five plants were used for either mock inoculation or *B. cinerea* infection, each plant representing a single biological replicate. The isolated total RNA (0.3 μg) was used to prepare samples using TruSeq^®^ Stranded mRNA (Illumina, San Diego, CA, USA) according to the manufacturer’s instructions. Briefly, total RNA was used to purify polyA containing mRNA molecules using poly-T oligo-attached magnetic beads. First strand and second strand cDNA were synthesized to obtain blunt ended ds cDNA. The 3□- ends of the blunt fragments were single adenylated and indexing adapters were ligated. The adapter-ligated fragments were selectively enriched by PCR amplification. The sample library was then validated by quantification, quality control and proceeded for normalisation and pooling of libraries. Library preparation and sequencing was performed at the Norwegian Sequencing Centre (Oslo, Norway) on the Illumina NextSeq 500 system with 75 bp high output single reads. The demultiplexed reads were aligned with the latest version of the *F. vesca* transcriptome (Edger *et al*., 2019) (ftp://ftp.bioinfo.wsu.edu/species/Fragaria_vesca/Fvesca-genome.v4.0.a1) using the aligner rsem (version 1.3.0) (Li and Dewey, 2011) with the bowtie2 option. Fragments per kilobase of exon per million reads mapped (FPKM) values were extracted from the gene results files and analyzed for differentially expressed genes using the R package EdgeR (Robinson, McCarthy and Smyth, 2009; McCarthy, Chen and Smyth, 2012). Normalization was performed using the trimmed mean of M values (TMM) method and p-values were adjusted using the Benjamini & Hochberg approach (Benjamini and Hochberg, 1995).

### 5.8 Transcriptome comparisons

We compared our RNA-seq data with previously published RNA-seq datasets on *F. vesca* responses to infection with *Phytophthora cactorum* (Toljamo *et al*., 2016) and *Colletotrichum acutatum* (Amil-Ruiz *et al*., 2016) and with datasets on *A. thaliana* responses to *B. cinerea* infection (Ferrari *et al*., 2007). Gene IDs from the published *F. vesca* studies (NCBI and Phytozome) were first matched with our data to obtain common gene IDs that were used to identify log_2_ fold-change (FC) values between studies. Corresponding homologs between *F. vesca* and *A. thaliana* were obtained by using the homology data from https://www.rosaceae.org/ (filed under: Fragaria_vesca/Fvesca-genome.v4.0.a1/homology/, file name: blastx_Fragaria_vesca_v4.0.a1_vs_TAIR10). The gene lists were filtered to retrieve the log_2_ FC values of differentially expressed genes (DEGs) with false discovery rate of less than 0.05 (FDR<0.05). The log_2_ FC values of these DEGs were used to create scatter plots and determine correlations between *F. vesca* responses to infection with two different pathogens and between *F. vesca* and *A. thaliana* responses to infection with *B. cinerea*. DEGs from different studies with FDR < 0.05 was used to generate Venn diagrams using InteractiVenn (Heberle *et al*., 2015).

### 5.9 Relative Growth Rate (RGR) assay

Young petioles on 8- to 12-week-old *F. vesca* plants were marked with plastic tags and petiole length was measured before and after treatments with priming chemicals or fungal infection. The following formula was used to calculate RGR: RGR = (ln h_2_ - ln h_1_)/t_2_ - t_1_, where ln = the natural logarithm, h2 and h_1_ = petiole length (in cm) at time t_2_ and t_1_ respectively (Buswell et al., 2018).

## 6. Acknowledgements

*Botrytis cinerea* Bc101 was provided by Gunn Mari Strømeng, NIBIO. We thank Jurriaan Ton for useful discussions and advice. This work was funded by a Toppforsk grant (249958/F20) from the Norwegian Research Council.

## 7. Author Contributions

RB, PK, TTh, MBB and CGF designed the experiments. RB and YZ performed the experiments. RB cultivated the plants for the experiments. RB wrote the first draft of the manuscript. TTe and RB analysed the RNA-seq data. TH provided plant material and greenhouse space for the imidacloprid experiment and demonstrated the method for plant growth measurements. RB, PK, TTh, MBB and CGF participated in regular discussions and work progress.

---

**Figure S1:**
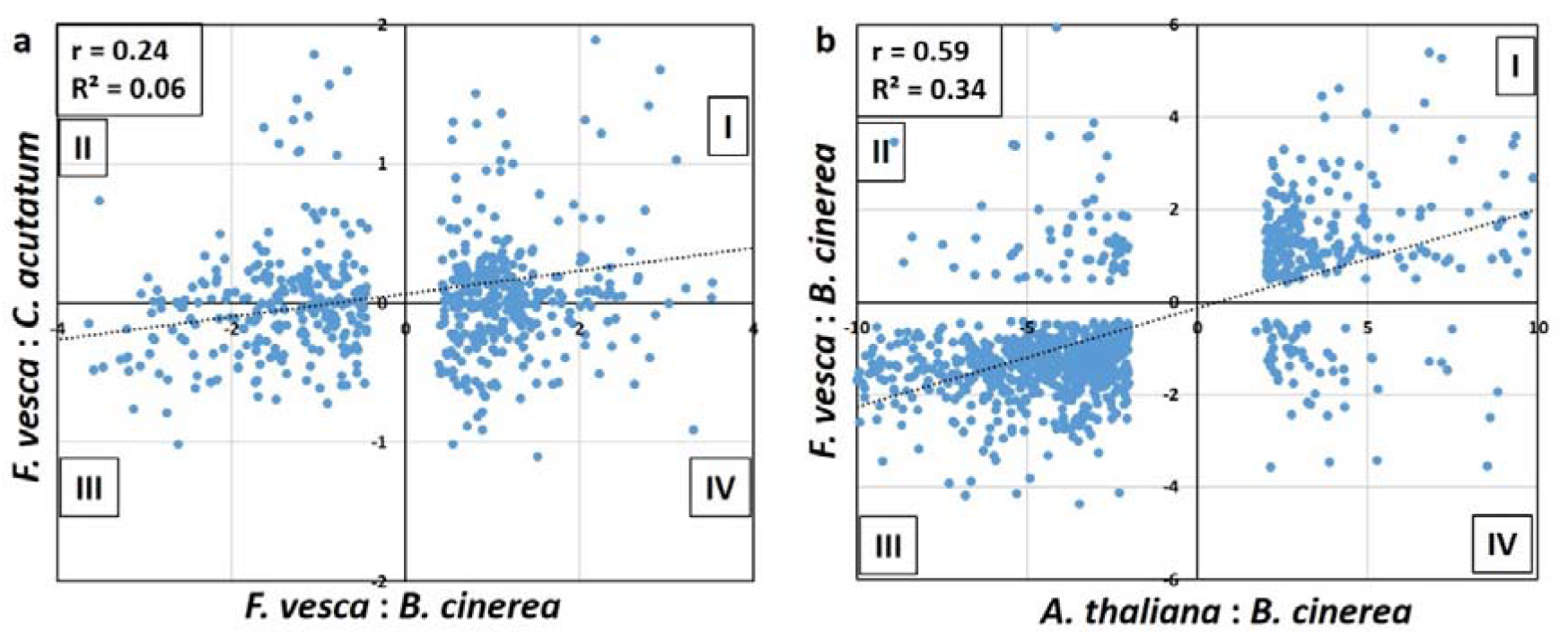
Scatter plots of log2 fold change values of differentially expressed genes (DEGs) common between (a) *Fragaria vesca* infected with *Botrytis cinerea* and *Colletotrichum acutatum* and (b) *Fragaria vesca* and *Arabidopsis thaliana, both infected with Botrytis cinerea*. The dots in quadrants I and III represent genes that respond similarly to both the pathogens, while the dots in II and IV quadrants represent the genes that respond oppositely to both pathogens. ‘r’ represents the correlation coefficient and ‘R^2^’ the coefficient of determination.

**Figure S2:**
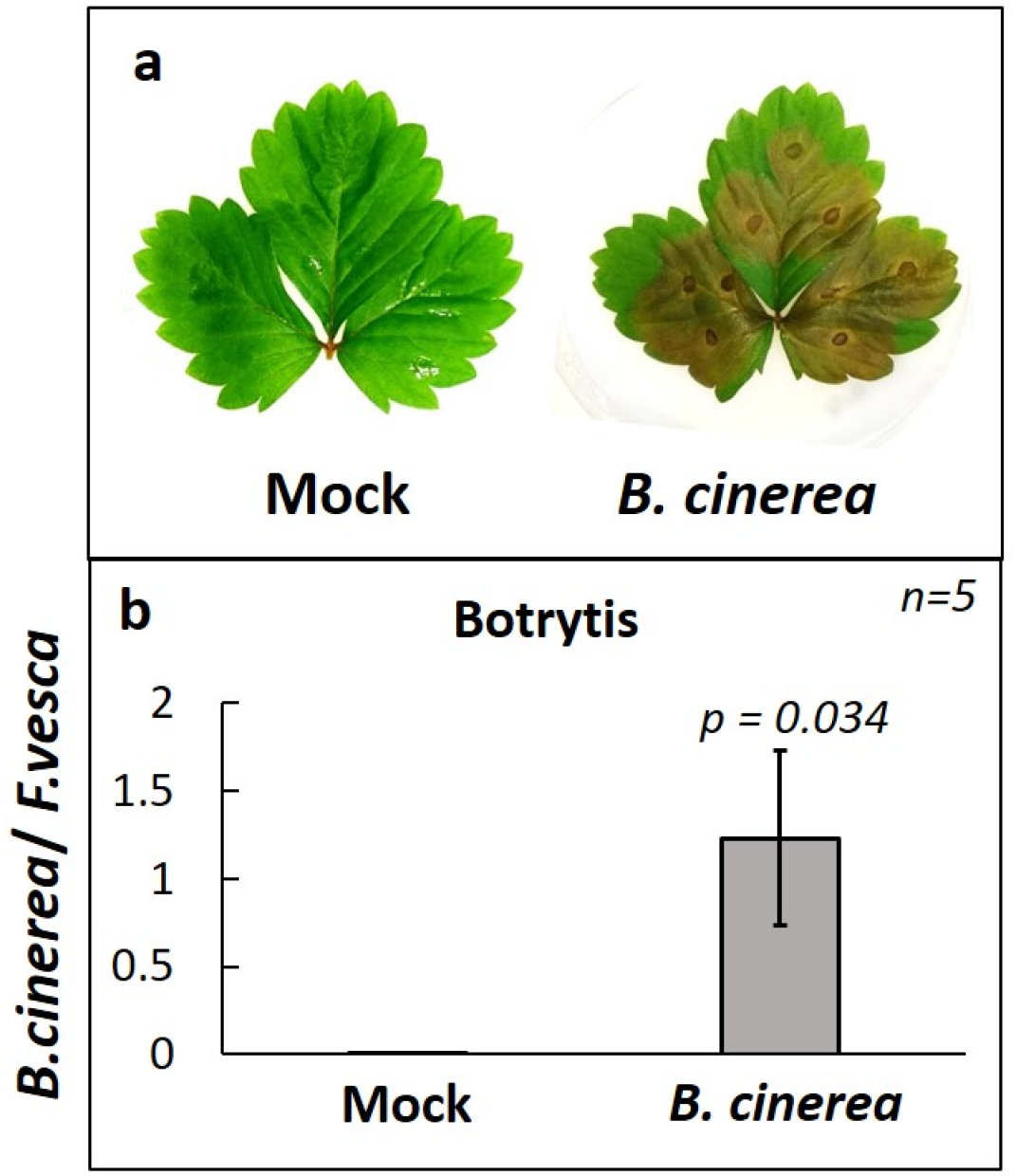
*Botrytis cinerea* infection of *Fragaria vesca* leaves. (a) Representative *Fragaria vesca* leaves showing infection by *Botrytis cinerea*, five days after drop inoculation of spores or mock suspension. (b) Genomic DNA-based qPCR quantification of relative amounts of Botrytis accumulation in infected and non-infected samples. ‘*n*’ represents number of biological replicates. Error bars represent standard error. Statistical significance was tested using student’s t-test.

**Figure S3:**
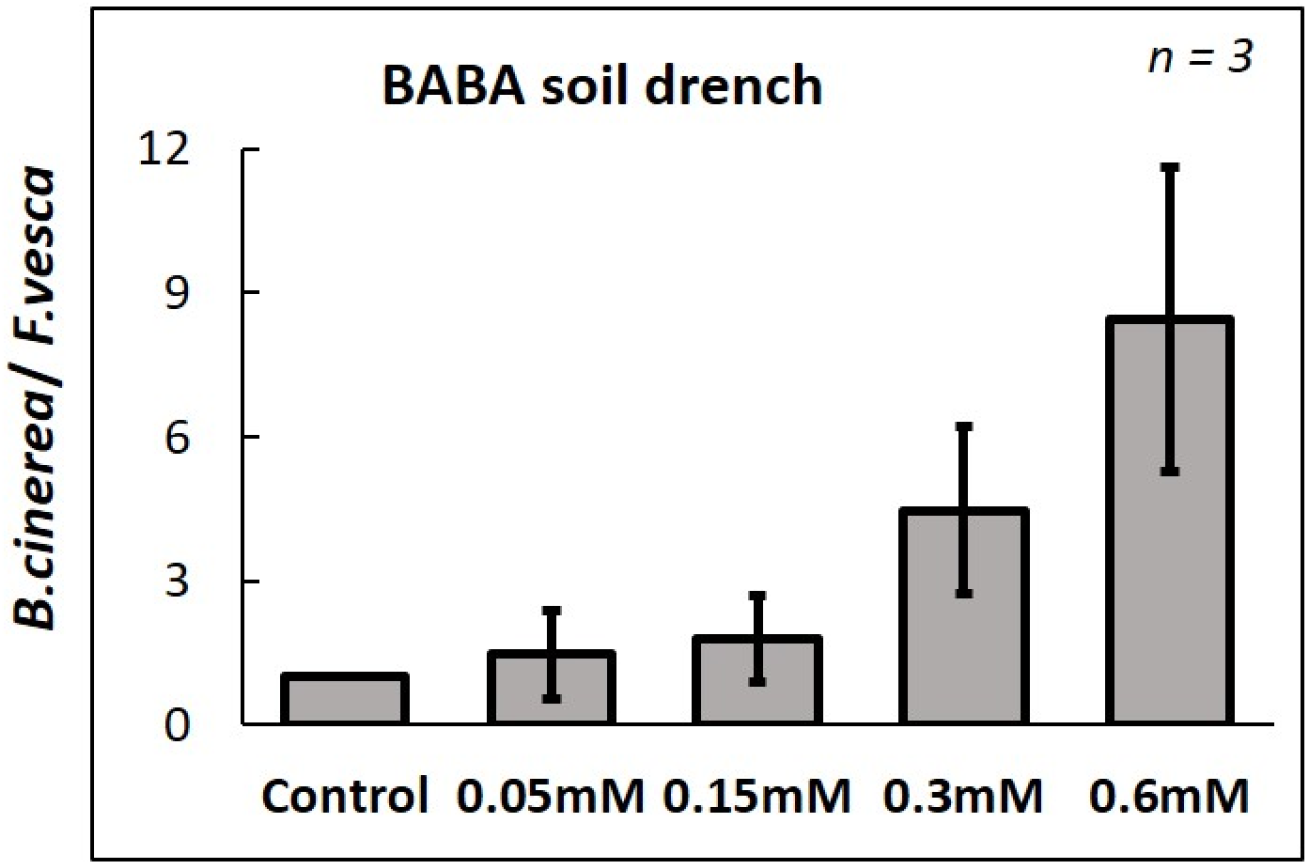
Effects of different concentrations of β-aminobutyric acid (BABA) on priming for resistance against *Botrytis cinerea* in *Fragaria vesca* ‘Alexandria’. Bars show amount of *Botrytis cinerea*, based on qPCR quantification, in plants soil-drenched with increasing concentrations of β-aminobutyric acid (BABA) and subsequently inoculated with *B. cinerea*. ‘*n*’ represents number of biological replicates. Error bars represent standard error.

**Figure S4:**
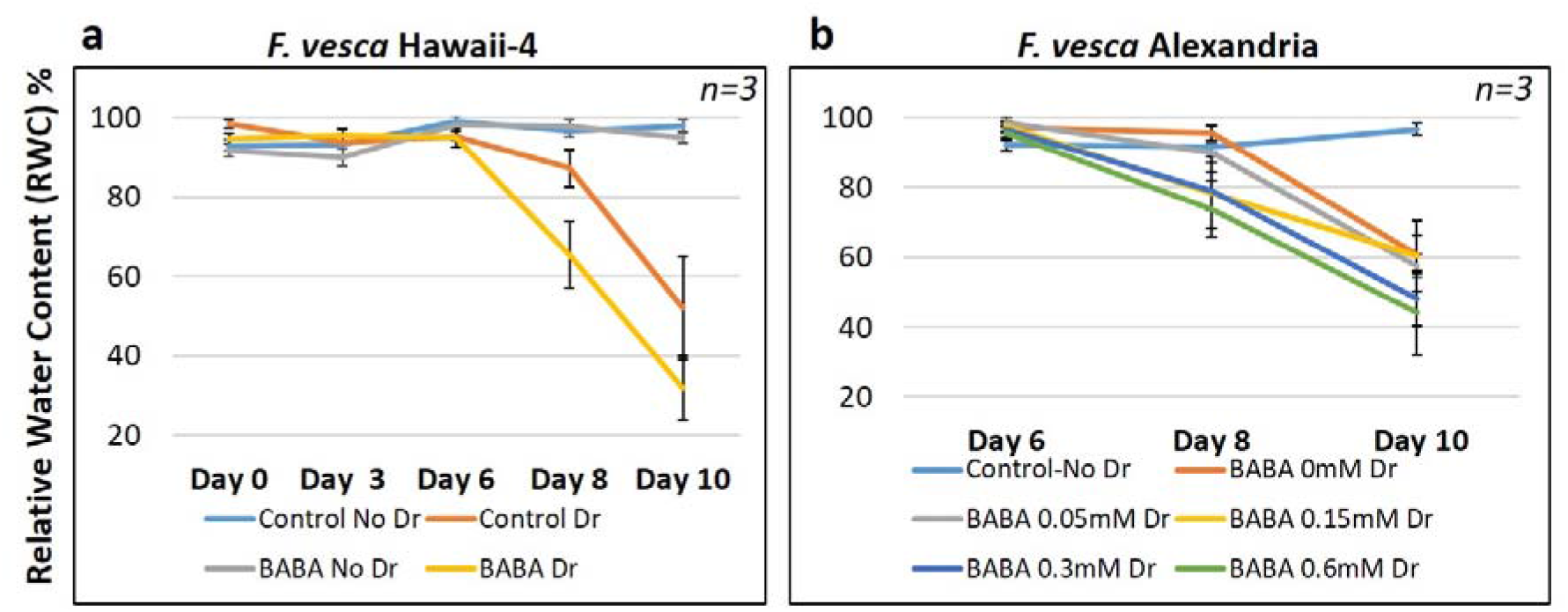
Effect of BABA soil-drench treatment on drought stress in *Fragaria vesca*. Relative water content (RWC) in plants soil-drenched with β-aminobutyric acid (BABA) in *F. vesca* Hawaii-4 (a) and in increasing concentrations of BABA in *F. vesca* Alexandria genotypes (b). ‘*n*’ represents number of biological replicates. Error bars represent standard error.

**Figure S5:**
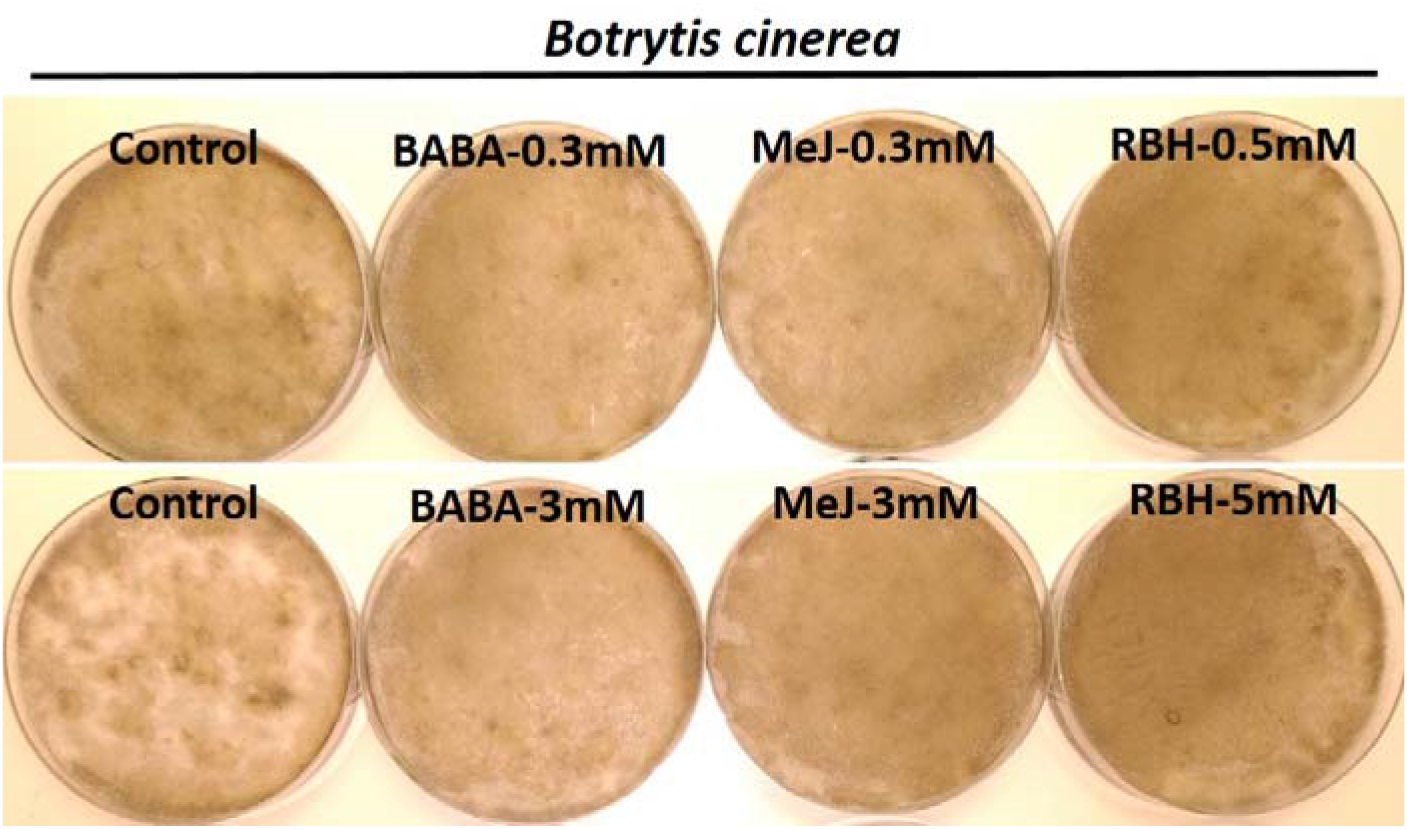
Growth of *Botrytis cinerea* (brown mycelium) on potato dextrose agar amended with different concentrations of the priming chemicals β-aminobutyric acid (BABA), methyl jasmonate (MeJ) and (*R*)-β-homoserine (RBH). Botrytis spore suspension was spread on the plates and incubated in dark for 4 weeks at room temperature before capturing photographs.

**Figure S6:**
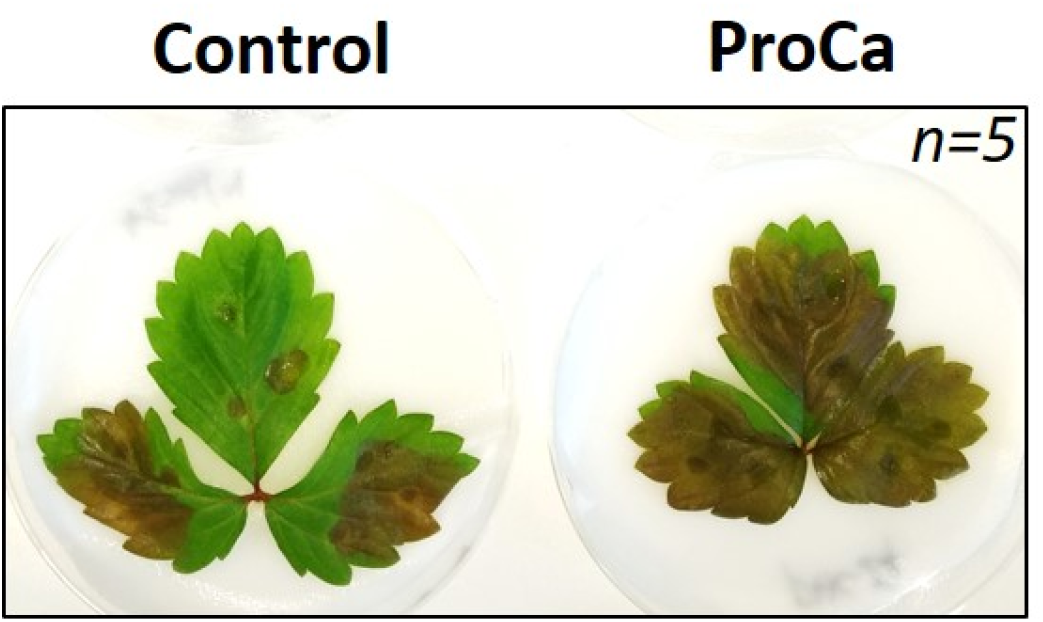
*Botrytis cinerea* induced necrotic lesions on *Fragaria vesca* leaves from plants sprayed two times with either Prohexadione-calcium (PHC) or control solution.

**Figure S7:**
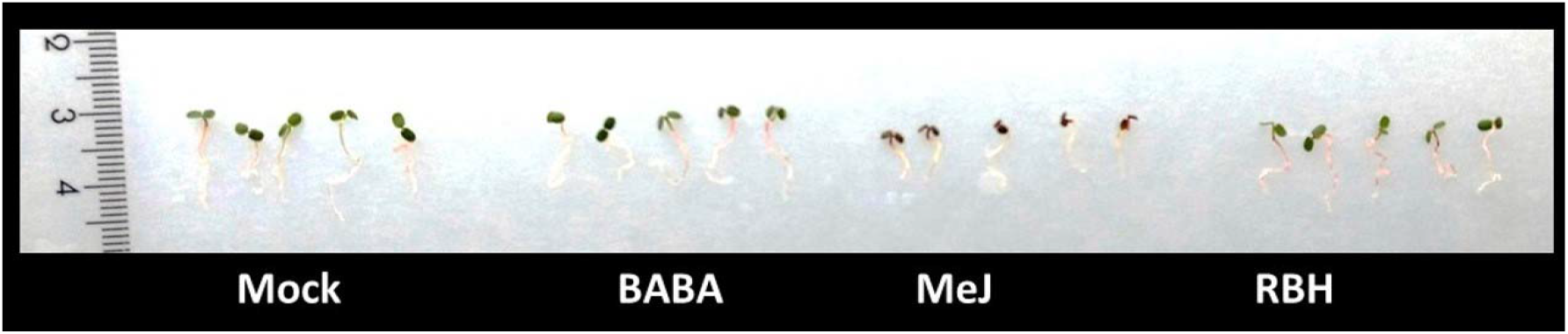
Representative 12-day-old *Fragaria vesca* seedlings germinated from seeds treated with the priming chemicals β-aminobutyric acid (BABA) (0.3mM), methyl jasmonate (MeJ) (0.3 mM) and (*R*)-β-homoserine (RBH) (0.5mM).

